# Dynamic Spiral Artery Remodeling in Early Human Pregnancy: An Analysis of Specimens Collected with a Standardized Protocol

**DOI:** 10.1101/2025.10.15.682615

**Authors:** Shenglong Ye, Yeling Ma, Wenlong Li, Linjing Qi, Xiao Fang, Xin Yu, Duo Yu, Xiaoye Wang, Dong-bao Chen, Yan-Ling Wang

## Abstract

Human uterine spiral artery remodeling (SAR) is a tightly regulated process involving complex interactions between interstitial and endovascular extravillous trophoblasts (iEVTs and enEVTs) and diverse maternal decidual cell populations. However, the intrinsic spatiotemporal dynamics of SAR in human placentation remain poorly understood, largely due to the limited availability of high-quality maternal-fetal interface specimens. Electively terminated early pregnancies offer a valuable resource for studying SAR *in situ*, yet inconsistent methods for distinguishing fragmented villous and decidual tissues have hindered reproducibility and interpretation. Herein we present a standardized protocol for the classification and characterization of high-quality maternal-fetal interface specimens from elective terminations by integrating stereomicroscopic evaluation with confirmation by immunohistochemistry and immunofluorescence microscopy. Combined with multiplex immunofluorescence imaging with cell-type– specific markers, this approach enabled precise spatial mapping and quantification of key morphological and cellular events in SAR from gestational weeks 5 to 10. Our analyses reveal that SAR initiates as early as week 5 with extraluminal recruitment of natural killer (NK) cells, followed by the formation of tightly packed EVT plugs within the lumens of spiral arteries in the decidua compacta; these plugs progressively extend deeper into the vessels and gradually loosen as gestation progresses. Notably, enEVTs appear to acquire NK cell-like phenotypes that may facilitate the displacement of endothelial and smooth muscle cells, promoting progressive vessel dilation. In summary, we provide a robust and reproducible method for assessing physiological SAR in early human pregnancy, supporting the adoption of our methodology in future studies of pathological SAR and related pregnancy disorders.

**Summary Sentence:** A standardized protocol for classifying maternal-fetal interface specimens from electively aborted materials at gestational weeks 5-10 was developed to quantify the spatiotemporal spiral artery remodeling dynamics in early human pregnancy.

**Graphic Abstract:** 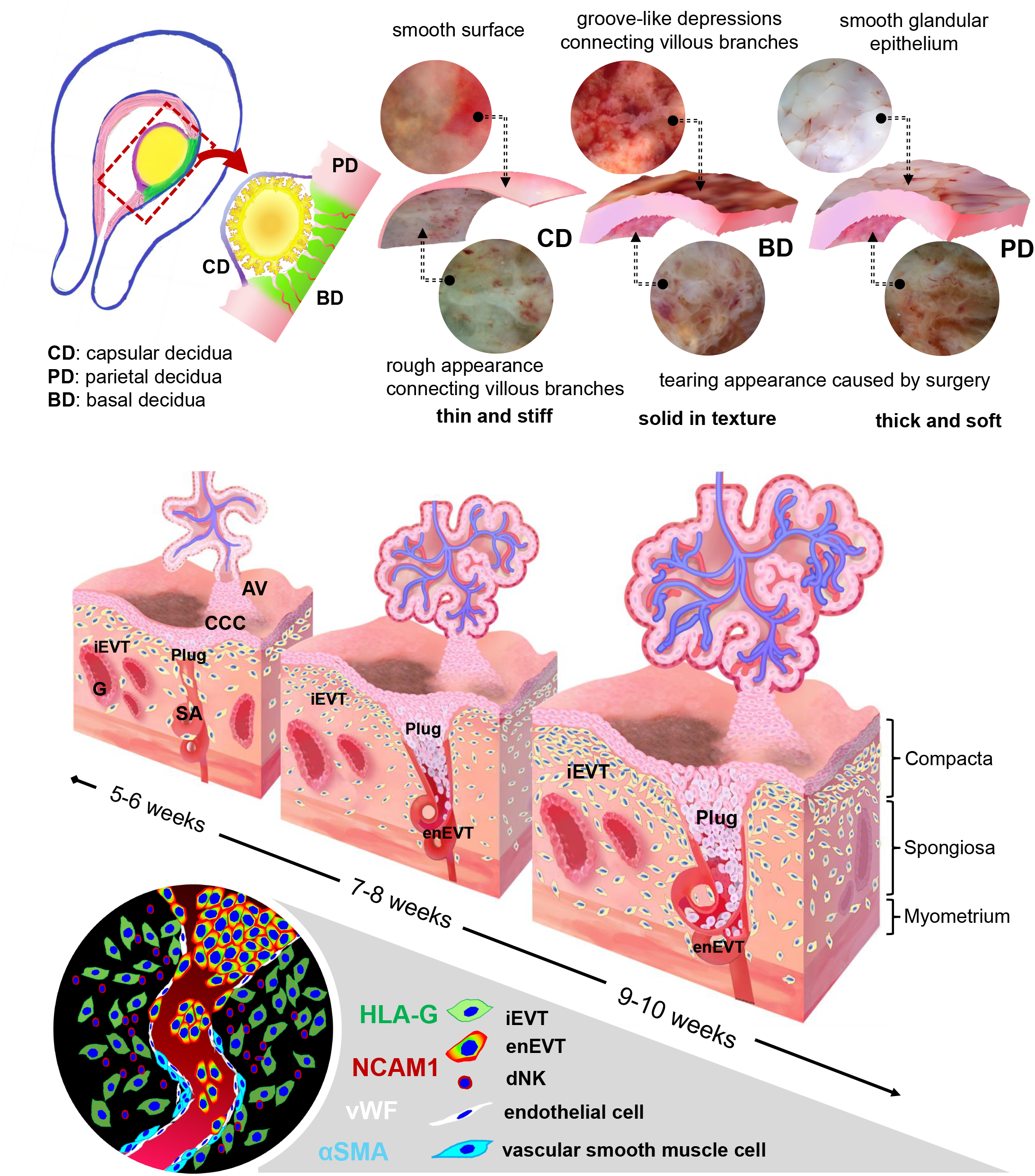

## INTRODUCTION

Human placentation begins approximately 5-7 days post-fertilization, when trophectoderm-derived extraembryonic cells first establish contact with the uterine endometrial lining. This process is initiated at the implantation site by the formation of the primitive syncytium, a multinucleated layer that encases the underlying mononuclear cytotrophoblast (CTB) cells. The primitive syncytium secretes proteolytic enzymes that degrade and loosen the surrounding decidual tissue, enabling its expansion into the resulting interstitial spaces [1]. By approximately two weeks post-fertilization (corresponding to ∼week 4 of gestation, counting from the last menstrual period), CTBs begin to migrate into these spaces and, together with mesenchymal cells, form the primitive placental villi. Continued proliferation and differentiation of CTBs give rise to the foundational villous architecture of the placenta, which consists of a mesenchymal core containing fetal blood vessels, an inner layer of villous CTBs, and an outer covering of syncytiotrophoblast (STB) [2]. Around gestational week 5, a subset of CTBs breaches the STB layer and differentiates into invasive extravillous trophoblasts (EVTs) [3-6]. These EVTs infiltrate the decidualized uterine stroma and extend into the superficial myometrium, facilitating physical anchorage of the developing placenta to the uterine wall.

The paired uterine arteries arise from the internal iliac arteries and give rise to a hierarchical vascular network comprising arcuate, radial, basal, and spiral arteries (SAs). Within the myometrium, the uterine artery branches into arcuate arteries, which course circumferentially and further subdivide into radial arteries. As the radial arteries approach the endometrial–myometrial junction, they give rise to basal arteries and SAs, the latter of which penetrate the endometrium. SAs are distinguished by their coiled morphology and serve as the primary blood supply to the functional layer of the endometrium [7]. During pregnancy, extravillous trophoblasts (EVTs) invade and form cell aggregates referred to as EVT plugs in the lumens of SAs. This invasion is accompanied by endothelial cell (EC) apoptosis, dedifferentiation of vascular smooth muscle cells (VSMCs), degradation of the extracellular matrix with fibrinoid deposition, and EVT incorporation into the vessel wall as endovascular EVTs (enEVTs) [8]. Collectively, these changes define the process of SA remodeling (SAR) which transforms the small-caliber, high-resistance SAs into dilated, low-resistance vessels with widened funnel-shaped openings into the intervillous space. This transformation enables the establishment of low-velocity maternal blood flow to minimize shear stress and to optimize placental perfusion that carries out the bi-directional maternal-fetal exchanges [9]. SAR occurs in coordination with decidualization and trophoblast differentiation/invasion processes, involving complex interactions between various fetal trophoblasts and maternal decidual, myometrial, immune, and vascular ECs and VSMCs. These events culminate in the formation of the maternal–fetal interface which is believed to structurally complete by gestational weeks 10–12 [10, 11]. Importantly, SAR is a critical event in placental development that underpins a successful pregnancy. Deficient or aberrant SAR is strongly associated with a spectrum of pregnancy complications, including early pregnancy loss, miscarriage, placental abruption, preeclampsia, and fetal growth restriction [12].

Despite extensive investigations, key questions remain regarding the intrinsic spatiotemporal dynamics of SAR. Ongoing debates concern the timing of SAR initiation [13, 14] and whether SAR occurs primarily intraluminally via enEVTs or extraluminally via interstitial EVTs (iEVTs) or other maternal cell types [11, 15, 16]. The mechanisms controlling the formation and resolution of EVT plugs also remain poorly understood, despite consistent evidence of their presence during varying gestation ages [17-20]. Using fresh maternal-fetal interface specimens, we recently observed EVT plugs within SAs that retained intact or partially disrupted vessel walls during early gestation [21, 22]. Complementing these findings, a digital quantification study of serial sections from the historical Boyd and Dixon placental collections demonstrated that EVT plugs persist within SA lumens throughout the first and second trimesters. These plugs become increasingly compact with advancing gestation, while their intercellular channel sizes expand [23]. These observations underscore persistent gaps in the understanding of EVT plug dynamics and associated cellular compositions. A more refined characterization of the spatiotemporal behavior of EVTs and associated maternal cells is needed to fully elucidate physiological SAR during early pregnancy.

A common denominator across many studies of SAR is the reliance on specimens sourced either from archived collections such as the Boyd collection or from fresh tissues obtained through elective abortions. However, variable protocols for sample collection, preservation, and processing may contribute to inconsistencies in SAR-related findings across different studies. More recently, archived hysterectomy-derived samples have been utilized to access intact maternal-fetal interface tissues for spatial transcriptomic analysis of SAR [24]. While valuable and cutting edge, such samples are extremely limited in availability, and consistent sampling across gestational stages remains a major challenge, resulting in spatial and temporal gaps in the understanding of SAR.

Elective abortion performed between gestational weeks 5–10 is legally permitted in certain countries, including China under strict regulations. This provides a unique opportunity to collect fresh villous and decidual tissues during a critical window of early placental development. However, these tissues are often fragmented by mechanical forces during surgical extraction, making it difficult to discern the native spatial architecture of the villous and decidual compartments. To overcome this challenge, we developed a standardized protocol for classifying maternal-fetal interface specimens obtained from the surgically removed materials during gestation weeks 5-10. By integrating multiplex immunofluorescence imaging with antibodies targeting specific cell markers at the maternal-fetal interface, our approach enables accurate delineation of spatial relationships among fetal trophoblasts and maternal decidual cells and vascular ECs and SMCs during weeks 5-10 of gestation. This methodology not only facilitates the reconstruction of the maternal-fetal interface but also provides novel insights into the spatiotemporal dynamics of SAR during early human placentation.

## MATERIALS and METHODS

### Ethics, human subjects, and sample collection

This study was approved by the Ethics Committees of the Institute of Zoology, Chinese Academy of Sciences (IOZ-2022-043), and the Peking University Third Hospital (2022 No. 379-02). Pregnant women who conceived naturally and opted for elective surgical abortion were recruited from the Family Planning Clinic at the Peking University Third Hospital (Beijing, China). Exclusion criteria included conception via assisted reproductive technologies, fetal chromosomal or congenital abnormalities, a history of renal or cardiovascular disease, and any gestational complications such as recurrent spontaneous miscarriage, gestational diabetes, hypertensive disorders of pregnancy, or intrauterine fetal death. Written informed consent was obtained for all tissue donations. Samples were collected from fifteen surgical procedures. Gestational age was determined by ultrasound measurement of embryo diameter immediately prior to surgery. Post-surgical materials, including villous and decidual tissues, were collected immediately, placed in ice-cold phosphate-buffered saline (PBS), transported to the research laboratory, and processed within one hour. The clinical characteristics of the study participants are summarized in **Table 1**.

**Table 1:** Clinical characteristics of participants.

| Case NO. | Gestational age<br>(Week<sup>+</sup>day) | Maternal age<br>(Year) | Parity<br>(Time) | Gravida<br>(Time) | BMI*<br>(Kg/m<sup>2</sup>) | Blood Pressure<br>(mm Hg. S/D)* |
| --- | --- | --- | --- | --- | --- | --- |
| 1 | 5<sup>+1</sup> | 31 | 1 | 1 | 24.84 | 115/69 |
| 2 | 6 | 38 | 0 | 1 | 22.03 | 103/55 |
| 3 | 6<sup>+3</sup> | 38 | 2 | 2 | 20.66 | 107/69 |
| 4 | 6<sup>+5</sup> | 39 | 2 | 2 | 22.04 | 115/61 |
| 5 | 6<sup>+5</sup> | 33 | 2 | 2 | 23.51 | 110/64 |
| 6 | 7<sup>+1</sup> | 38 | 2 | 3 | 25.21 | 114/64 |
| 7 | 7<sup>+5</sup> | 27 | 0 | 0 | 18.37 | 97/62 |
| 8 | 7<sup>+6</sup> | 25 | 1 | 2 | 18.17 | 103/65 |
| 9 | 8 | 35 | 0 | 2 | 18.03 | 126/68 |
| 10 | 8<sup>+2</sup> | 30 | 0 | 0 | 18.78 | 125/65 |
| 11 | 9<sup>+2</sup> | 38 | 0 | 0 | 20.25 | 107/66 |
| 12 | 9<sup>+2</sup> | 38 | 2 | 2 | 20.31 | 102/61 |
| 13 | 9<sup>+4</sup> | 21 | 0 | 0 | 19.63 | 106/67 |
| 14 | 9<sup>+6</sup> | 31 | 1 | 2 | 20.20 | 111/64 |
| 15 | 10<sup>+1</sup> | 33 | 0 | 0 | 17.92 | 115/63 |

BMI: Body mass index; S/D: Systolic/Diastolic blood pressure.

### Stereomicroscopic separation of decidua and placenta villi

Aborted tissues were rinsed with ice-cold PBS to remove blood clots and then immersed in fresh PBS. Under a 50x stereomicroscope, placental villi were identified based on the morphological characteristics of the chorionic sac. Villi appeared as white, floating tissues with a densely branched structure. Remaining tissues were separated and identified as decidua, comprising various decidual subtypes (**Fig. 1A**). Decidual tissues were further classified into three types based on macroscopic features, including thickness, stiffness, surface smoothness, and the presence or absence of residual villous tissues. These classifications were subsequently validated by immunohistochemical analysis.

**Figure 1.**
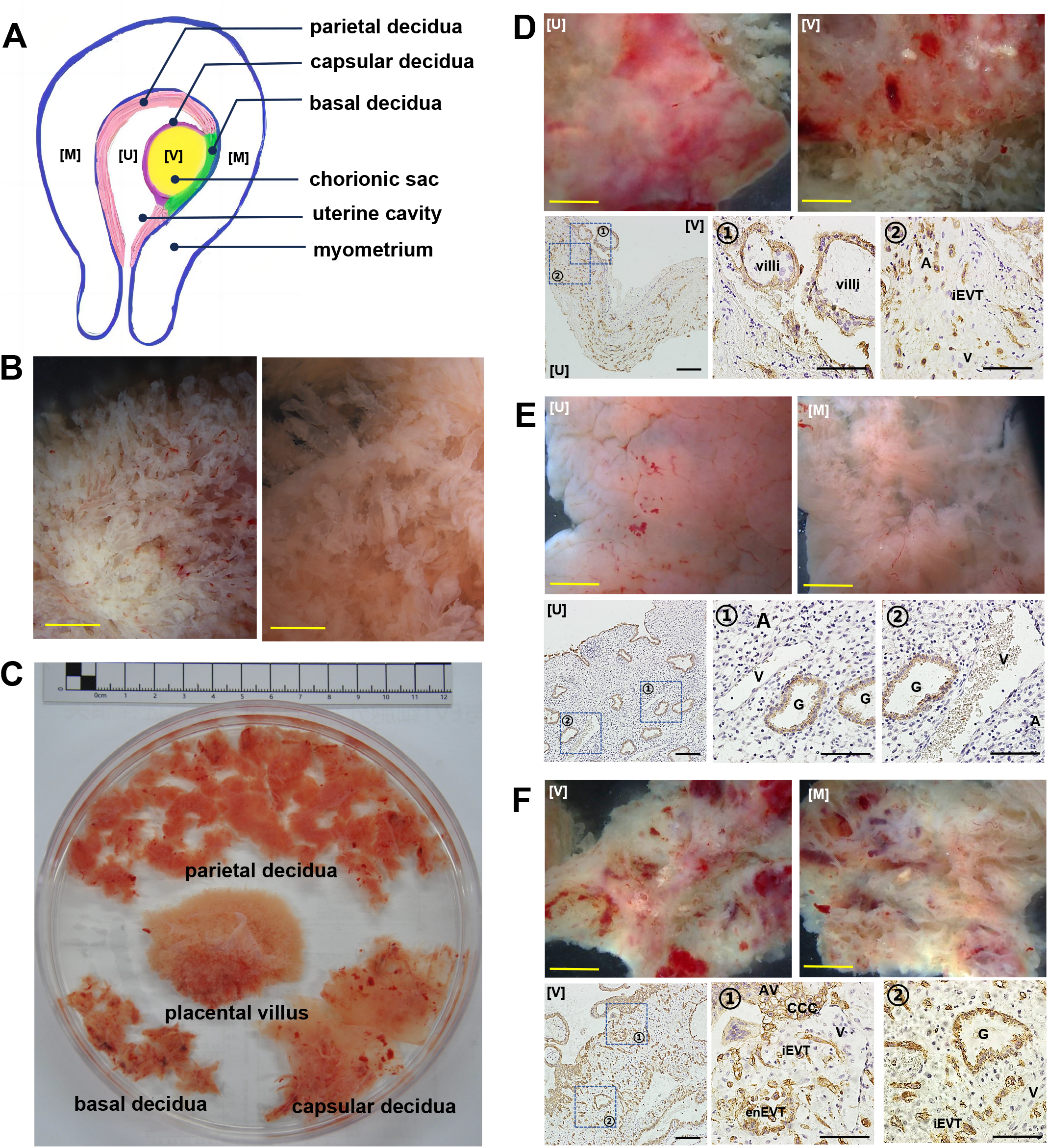
Classification of human early pregnancy abortion tissues. A. Schematic diagram showing the chorionic sac and the three types of decidual tissues: decidua basalis, capsularis, and parietalis. B. Representative gross morphology of villous tissues at 6 weeks (right) and 9 weeks (left) of gestation, imaged under a stereomicroscope at 50× magnification. C. Gross morphology of aborted tissues at 8 weeks of gestation, classified into chorionic sac, decidua parietalis, decidua basalis, and decidua capsularis. D. Gross morphology (top) of decidua capsularis at the uterine luminal side [U] and villous side [V]; corresponding immunohistochemical staining for cytokeratin-7 (CK7, bottom) to visualize trophoblasts. E. Gross morphology (top) of decidua parietalis at the uterine luminal side [U] and the myometrial detachment surface [M]; CK7 immunostaining (bottom) highlights the absence of trophoblasts. F. Gross morphology (top) of decidua basalis at the villous side [V] and muscle layer detachment surface [M]; CK7 immunostaining (bottom) shows abundant trophoblast infiltration. G, gland; iEVT, interstitial extravillous trophoblast; enEVT, endovascular extravillous trophoblast; SA, spiral artery; V, vein; AV, anchoring villi; CCC, cytotrophoblastic cell column. Scale bars: Black = 50 μm; Yellow = 2 mm.

### Paraffin-embedded section preparation

The separated villous and decidual tissues were immediately fixed in 4% formaldehyde at 4 °C for 12 hours. After fixation, samples were rinsed with PBS, dehydrated through a graded ethanol series, and cleared in xylene. For basal decidual tissues, special care was taken during paraffin embedding to orient the specimens such that serial sections extended from the superficial implantation layer (decidua compacta) to the deeper layer (decidua spongiosa). Paraffin blocks were sectioned at 5 μm thickness; the sections were stored at room temperature (RT) until further analysis.

### Immunohistochemistry

Paraffin sections were dewaxed in xylene and rehydrated through a graded ethanol series. Endogenous peroxidase activity was quenched using hydrogen peroxide. Antigen retrieval was achieved by heating the sections in trisodium citrate dihydrate buffer. Following antigen retrieval, nonspecific binding was blocked with 3% fetal bovine serum albumin (FBS) in PBS. Sections were then incubated overnight at 4 °C with a primary antibody against human cytokeratin 7 (CK7, 1:8000; Abcam #ab181598). After three washes (5 minutes each) in PBS, sections were incubated with horseradish peroxidase (HRP)-conjugated secondary antibodies (PV-6001; ZSGB-BIO, China) for 1 hour at room temperature. Immunoreactivity was visualized using diaminobenzidine (DAB; ZLI-9019, ZSGB-BIO, China) as the chromogen, followed by hematoxylin counterstaining. Digital images were acquired using a light microscope equipped with a 10x ocular lens and 2x, 10x, and 20x air objectives, and captured with an Olympus DP72 CCD camera (Olympus, Japan).

### Multiplexed immunofluorescence imaging of SAR

Multiplexed immunofluorescence microscopy for imaging analysis of SAR was performed using the Opal multiple-color manual IHC kit (Akoya Biosciences, Marlborough, MA, USA), which includes Opal 520, 570, 620, 650, and 690 fluorophores. Each fluorophore was used to label HRP-conjugated secondary antibodies (Vector Laboratories, Burlingame, CA, USA), enabling the sequential detection of five distinct primary antibodies targeting different cellular markers within the same tissue section. Briefly, sections of maternal–fetal interface tissue were dewaxed in xylene and rehydrated through a graded ethanol series. After blocking endogenous peroxidase activity and antigen retrieval, sections were blocked with Opal blocking buffer at RT for 10 minutes, then incubated with a primary antibody diluted in Opal antibody diluent at 37 °C for 1 hour. After washing three times for 5 minutes each with Tris-buffered saline containing Tween-20 (TBST), sections were incubated with an HRP-conjugated secondary antibody at RT for 10 minutes. After another series of TBST washes, the appropriate Opal fluorophore was applied according to the manufacturer’s instructions at 37 °C for 6 minutes. This staining cycle was repeated sequentially for each of the five primary antibodies. Upon completion of all staining steps, sections were washed thoroughly and mounted using a DAPI-containing antifade mounting medium (ZSGB-BIO, ZLI-9557) for nuclear counterstaining. Imaging was performed using a Leica Stellaris 5 confocal microscope (Leica Microsystems, Germany) equipped with a 10**x** ocular lens and 20**x** air or 40**x** oil immersion objectives. Whole-section images were acquired for analysis of dynamic cellular and structural changes, including the presence of EVT plugs in SAs, using the LAS-X FLIM/FCS module (Leica Microsystems). The analysis included at least ten sections per sample, with sections taken from every 20th slice across a total of more than 200 slices continuously collected. Each fluorophore corresponded to a specific target protein, detected using white laser excitation at the following wavelengths (nm): 494 (Opal 520), 550 (Opal 570), 588 (Opal 620), 627 (Opal 650), 676 (Opal 690), and 368 (DAPI). Corresponding emission detection ranges were set to 497–544, 564–606, 617–630, 633–653, 682–762, and 420–467 nm, respectively.

The following primary antibodies were used for multiplex immunofluorescence staining: anti-cytokeratin 7 (CK7, 1:8000; Abcam, #ab181598), anti-human leukocyte antigen-G (HLA-G, 1:200; Abcam, #ab52455), anti-E-cadherin (E-cad, 1:200; Cell Signaling Technology, #3195S), anti-neural cell adhesion molecule 1 (NCAM1, 1:200; Abcam, #ab75813), anti-von Willebrand factor (vWF, 1:200; Abcam, #ab6994), and anti-α-smooth muscle actin (αSMA, 1:400; Abcam, #ab5694). CK7 was used as a general trophoblast marker[25], while HLA-G served as a specific marker for extravillous trophoblasts (EVTs) [26]. E-cadherin (E-cad), an epithelial cell marker, is frequently expressed during EVT differentiation [8]. NCAM1 was used to label decidual natural killer (dNK) cells [27]. Interstitial EVTs (iEVTs) were identified as HLA-G^+^/NCAM1^−^ cells, whereas endovascular EVTs (enEVTs) were defined as HLA-G^+^/NCAM1^+^ cells [28, 29]. vWF was used to identify endothelial cells (ECs) [30], and αSMA was used to mark vascular smooth muscle cells (VSMCs) [31].

### Spatiotemporal analysis of SAR

Basal decidua samples containing the maternal–fetal interface were collected from 15 elective abortion surgeries (**Table 1**), paraffin-embedded, and sectioned for spatiotemporal analysis of SAR across different gestational ages and decidual zones (compacta vs. spongiosa). From each sample, approximately 200 consecutive sections (5 μm) were cut. For analysis, every 20th section was selected, resulting in at least 10 sections per sample. Multiplex immunofluorescence microscopy was performed on these sections using antibodies against HLA-G, NCAM1, vWF, and αSMA to label extravillous trophoblasts (EVTs), decidual natural killer (dNK) cells, endothelial cells (ECs), and vascular smooth muscle cells (VSMCs), respectively. SAR was classified into three distinct stages based on cellular distribution and vascular structure, as previously described [32]:

- Unremodeled SAs - characterized by multiple layers of VSMCs and intact endothelium without EVT infiltration.
- Actively remodeling SAs - identified by EVT invasion beyond the vessel wall, partial disruption of the VSMC layer, and initiation of EC replacement.
- Fully remodeled SAs - marked by complete replacement of both ECs and VSMCs by EVTs.

### Statistical analysis

Data were expressed as the mean ± standard deviation (SD). Values were calculated either as percentages or as fold changes relative to gestational week 5. Statistical analyses were performed using GraphPad Prism version 9.0 (GraphPad Software, San Diego, CA, USA). Differences between two groups were assessed using Student’s *t*-test, while comparisons among multiple groups were conducted using one-way analysis of variance (ANOVA), followed by Bonferroni post hoc tests when appropriate. A *P*-value of less than 0.05 was considered statistically significant.

## RESULTS

### Tissue composition and characteristics of decidual subtypes

Elective abortion surgeries resulted in the complete removal of both decidual and fetal tissues [23], as confirmed in **Fig. 1A**. The recovered tissues included fetal components such as the chorionic sac and placental villi and maternal uterine decidua. The decidua comprises three morphologically distinct regions: 1) basal decidua located between the chorionic sac (V) and the uterine myometrium (M); 2) capsular decidua surrounding the chorionic sac (V) and facing the uterine cavity (U); and 3) parietal decidua positioned opposite the basal decidua, lining the remainder of the uterine cavity. Morphologically, fetal-derived placental villi displayed a densely branched, tree-like structure, with increasing complexity observed at later gestational stages (**Fig. 1B**). The collected specimens typically included 20–40 tissue fragments, which were grossly classified into basal, capsular, or parietal decidua based on surface characteristics and structural features (**Fig. 1C**).

Approximately 20% of the decidual fragments were thin and stiff, with thicknesses ranging from 0.05 to 0.1 mm (**Fig. 1C**). Stereomicroscopic examination revealed a smooth surface on one side and a rough surface on the other, with occasional attachment of villous branches (**Fig. 1D**, top). These features are characteristic of capsular decidua, which lies on the surface of the chorionic sac. As gestation progressed from weeks 5 to 10, capsular decidua specimens became progressively thinner, consistent with the expansion of the chorionic sac. Immunohistochemical staining for CK7 confirmed the presence of dispersed villi and EVTs, accompanied by sparse vasculature (**Fig. 1D**, bottom), hallmark histological features of capsular decidua.

Approximately 70% of the decidual specimens were soft tissues, measuring 1.0 to 3.0 mm in thickness (**Fig. 1C**). One side of these specimens exhibited a smooth, glandular epithelial surface, with underlying blood vessels clearly visible. Stereoscopic examination revealed a smooth surface with gland-like openings on one side, and a relatively rough opposite surface marked by clear evidence of connective tissue tearing likely resulting from surgical separation (**Fig. 1E**, top). Immunohistochemical staining for CK7 confirmed the presence of abundant glands and some intact blood vessels, but notably, a lack of EVTs within the stroma (**Fig. 1E**, bottom). These features are consistent with the parietal decidua.

Less than 10% of the decidual specimens appeared as solid tissues, with a thickness ranging from approximately 0.5 to 1.5 mm (**Fig. 1C**). Stereoscopic observation showed that both surfaces were rough. One side displayed numerous characteristic groove-like depressions, loose gland-like openings, and a relatively high density of residual villi; the opposite surface exhibited tearing features similar to those seen in parietal decidua (**Fig. 1F**, top). CK7 immunostaining revealed hallmark features of the basal decidua containing the maternal–fetal interface, including anchoring villi with cytotrophoblastic cell columns (CCCs) embedded in the decidual stroma, abundant iEVTs, EVT plugs, enEVTs within uterine blood vessels, and expanded glands lined by a thin epithelial layer (**Fig. 1F**, bottom).

### Identification of maternal-fetal interface in basal decidua

The maternal-fetal interface forms at the site of blastocyst implantation. Surgical abortion disrupts the structural integrity between placental villi and the decidua, highlighting the need to carefully identify and isolate intact maternal-fetal interface tissue from the mixture of removed tissues for studying physiological SAR during early human pregnancy. Using stereomicroscopy, basal decidual tissues containing residual villi embedded within or surrounding groove-like depressions were meticulously separated (**Fig. 2A**). CK7 staining of longitudinal tissue sections revealed the typical structural features of an intact maternal-fetal interface. Based on the spatial distribution of CK7-positive uterine glands, the decidua was categorized into two layers: compacta and spongiosa decidua (**Fig. 2B**), corresponding to superficial and deep regions of embryonic implantation, respectively [28].

**Figure 2.**
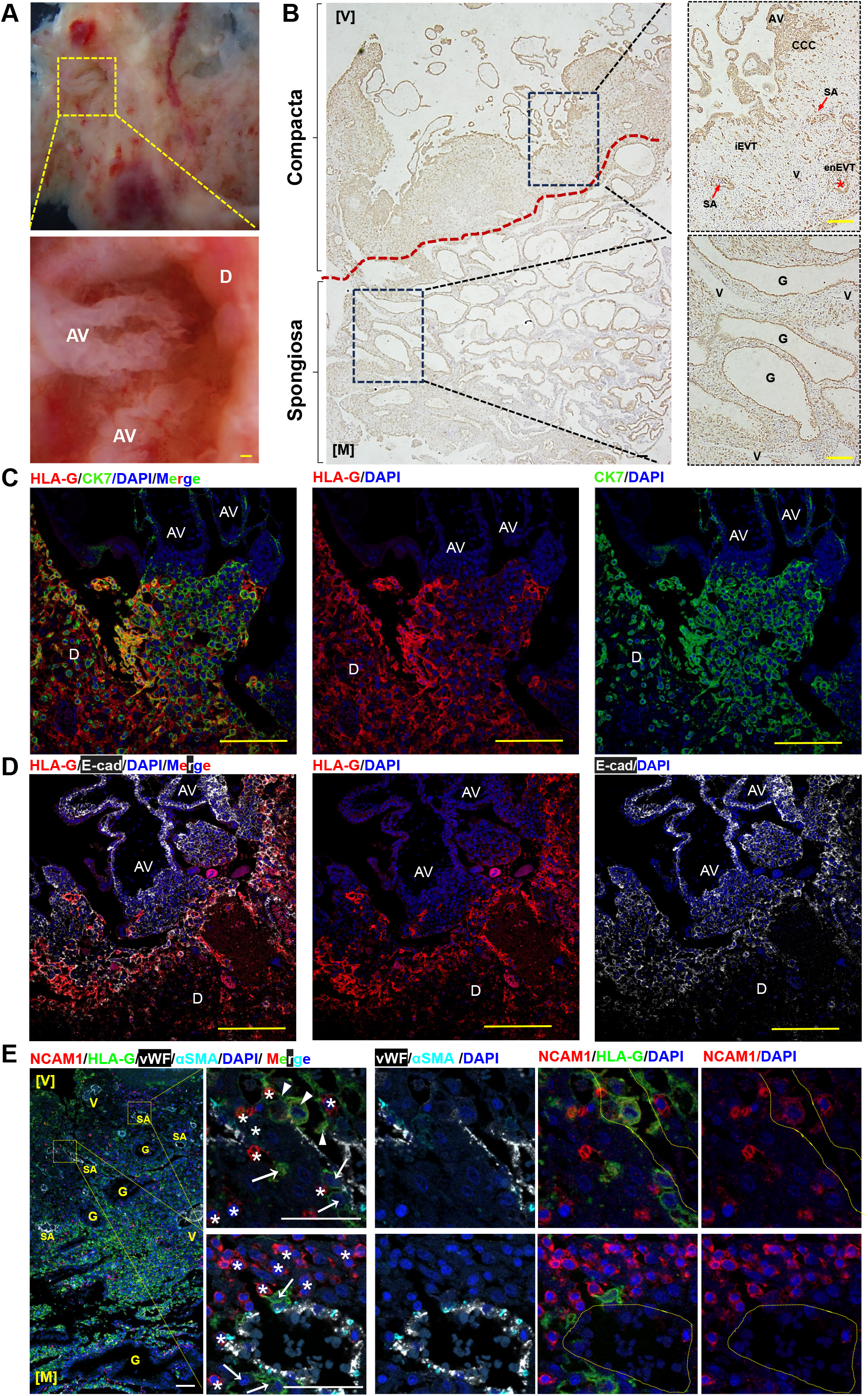
Morphological characteristics of human decidua basalis during 5–10 weeks of gestation. A. Gross morphology of a gestational week (GW) 9 decidua basalis imaged from the villous-facing side under 50× stereomicroscope, showing a characteristic pale depression structure where the villous tree anchors to the decidua basalis. B. Representative immunofluorescence staining of cytokeratin-7 (CK7) in a GW9 decidua basalis, showing distinct compacta and spongiosa layers separated by a red dashed line based on gland distribution. C. Representative immunofluorescence images of CK7 (green) and human leukocyte antigen G (HLA-G, red) in a GW7 decidua basalis with anchoring villi. Nuclei are counterstained with DAPI (blue). D. Representative immunofluorescence staining of HLA-G (red) and E-cadherin (white) in a GW7 decidua basalis with anchoring villi. DAPI marks nuclei (blue). E. Representative immunofluorescence image of a GW9 decidua basalis showing spiral arteries at different stages of remodeling. HLA-G (green) labels extravillous trophoblasts (EVTs), NCAM1 (red) marks uterine natural killer (uNK) cells, von Willebrand factor (vWF, white) labels endothelial cells (ECs), α-smooth muscle actin (αSMA, cyan) marks smooth muscle (SM) cells, and DAPI (blue) marks nuclei. uNK cells, iEVTs, and enEVTs are marked by star, arrow, and arrowhead, respectively. AV, anchoring villus; D, decidua; E-cad, E-cadherin; V, villous-facing side; M, myometrial-facing (uterine muscle detachment) side. Scale bars: Yellow = 100 μm; White = 50 μm.

Immunofluorescence microscopy identified CK7-positive anchoring villi and CK7/HLA-G double-positive trophoblasts derived from distal cytotrophoblast cell columns (CCCs) within the compacta layer (**Fig. 2C**). E-cadherin (E-cad) staining showed a progressive decrease in intensity from villous trophoblasts to interstitial EVTs (iEVTs) infiltrating the decidual stroma (**Fig. 2D**), suggesting epithelial-mesenchymal transition (EMT) during EVT differentiation, which is consistent with previous findings [33]. Multiplex immunofluorescence analyses were performed using markers for endothelial cells (vWF), vascular smooth muscle cells (αSMA), EVTs (HLA-G), iEVTs (HLA-G^+^/NCAM1^−^), and endovascular EVTs (enEVTs; HLA-G^+^/NCAM1^+^) to delineate spatial interactions among key fetal-derived (iEVT, enEVT) and maternal-derived (ECs, VSMCs, dNK cells) cell types at the maternal–fetal interface (**Fig. 2E**). Blood vessels positive for vWF and/or αSMA were observed in both the compacta and spongiosa layers. HLA-G^+^/NCAM1^+^ enEVTs were detected within SAs exhibiting partial or complete disruption of EC and VSMC layers, indicating various stages of SAR. Notably, the SMC layers of actively remodeling SAs were frequently infiltrated by NCAM1^+^ decidual natural killer (dNK) cells. These remodeling vessels showed significantly higher dNK cell clustering density during early gestation, suggesting a potential role for dNKs in the early phases of SAR.

### Spatiotemporal dynamics of SAR

Successful classification of intact maternal-fetal interface samples enables accurate analysis of spatiotemporal changes in SAR during early human development. We optimized a multiplexed immunofluorescence microscopy approach to analyze longitudinal sections of the maternal-fetal interface from 5 to 10 weeks of gestation. Consistent with our previous findings [32], un-remodeled SAs appeared small, characterized by an intact vWF-positive EC lining and multilayered αSMA-positive VSMCs. These vessels were surrounded with or without NCAM1^−^/HLA-G^+^ iEVTs, but critically, all lacked luminal NCAM1^+^/HLA-G^+^ enEVTs. In contrast, actively remodeling SAs showed enlarged lumens and were surrounded by NCAM1^−^/HLA-G^+^ iEVTs. The EC/VSMC layers were partially diminished and progressively replaced by NCAM1^+^/HLA-G^+^ enEVTs within the lumen. Fully remodeled SAs displayed the largest lumens, complete replacement of ECs by NCAM1^+^/HLA-G^+^ enEVTs, which were also surrounded by NCAM1^−^/HLA-G^+^ iEVTs (**Fig. 3A**).

**Figure 3.**
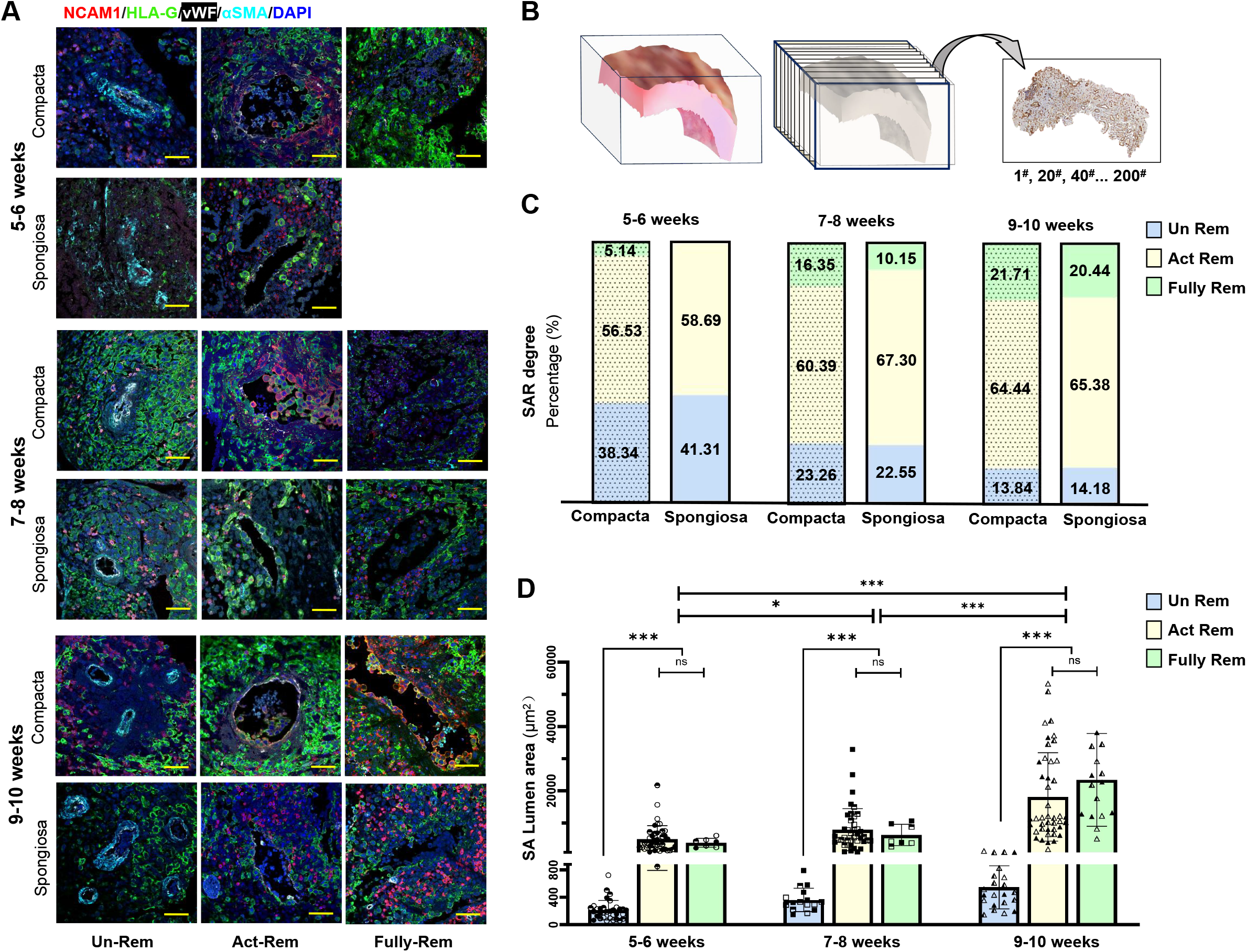
Dynamic spiral artery remodeling (SAR) in human decidua basalis during 5–10 weeks of gestation. A. Representative multiplex immunofluorescence images showing spiral arteries (SAs) in the compacta and spongiosa layers of the decidua basalis at different gestational stages. Green: HLA-G (extravillous trophoblasts); Red: NCAM1 (uterine natural killer [uNK] cells); White: von Willebrand factor (vWF, endothelial cells); Cyan: α-smooth muscle actin (αSMA, smooth muscle cells); Blue: DAPI (nuclei). SAs were categorized as: Un-remodeled (Un-Rem): intact αSMA-positive smooth muscle (SM) and vWF-positive endothelial cell (EC) layers (left panel); actively remodeling (Act-Rem): partially disrupted SM layer with infiltration of HLA-G^+^ EVTs and NCAM1^+^ uNK cells (middle panel); and fully remodeled (Fully-Rem): complete loss of SMC/EC layers replaced by HLA-G^+^ EVTs and NCAM1^+^ uNK cells (right panel). Scale bar: 50 μm. B. Schematic of sectioning strategy. Approximately 200 serial paraffin sections were collected per case; every 20th section was used for quantitative analysis of SAR. C. Bar graph showing the percentage of un-remodeled, actively remodeling, and fully remodeled SAs in compacta and spongiosa layers, stratified by gestational age. Detailed statistics are provided in Table 2. D. Bar graph summarizing luminal areas of SAs in different remodeling stages (Un-Rem, Act-Rem, Fully-Rem) at each gestational age. Each symbol represents an individual case (n=3 per group). All SAs within each analyzed section were counted. Pairwise comparisons were made with t-tests. *, *P* < 0.05; **, *P* < 0.01; ***, *P* < 0.001

We grouped 15 maternal-fetal interface specimens into three gestational age categories: 5–6 weeks, 7– 8 weeks, and 9–10 weeks (n=3 per group; **Table 2**). SAR dynamics were quantified based on gestational age and trophoblast invasion depth (decidua compacta vs. spongiosa) using 10 sections from every 20th slice from a total of 200 sections continuously cut from each case (**Fig. 3B**). The extent of SAR increased significantly from 5-6 to 9-10 weeks of gestation. Specifically, the proportion of un-remodeled SAs decreased from approximately 40% at 5–6 weeks to 14% at 9–10 weeks, while fully remodeled SAs increased from 5% to 20% during the same interval (**Fig. 3C, Table 2**). At 5-6 weeks, the spongiosa region exhibited a higher proportion of un-remodeled SAs compared to the compacta. However, by 7-8 and 9-10 weeks, SAR progression became comparable between these regions (**Fig. 3C, Table 2**). Lumen area measurements revealed a significant increase across gestation. The mean lumen areas of actively remodeling and fully remodeled SAs were: 4,840.79 ± 3,936.71 μm^2^ at 5-6 weeks, 7,709.08 ± 6,179.37 μm^2^ at 7-8 weeks, and 19,586.78 ± 13,991.63 μm^2^ at 9-10 weeks (**Fig. 3D**), with statistically significant differences among the groups (p < 0.001).

**Table 2:**
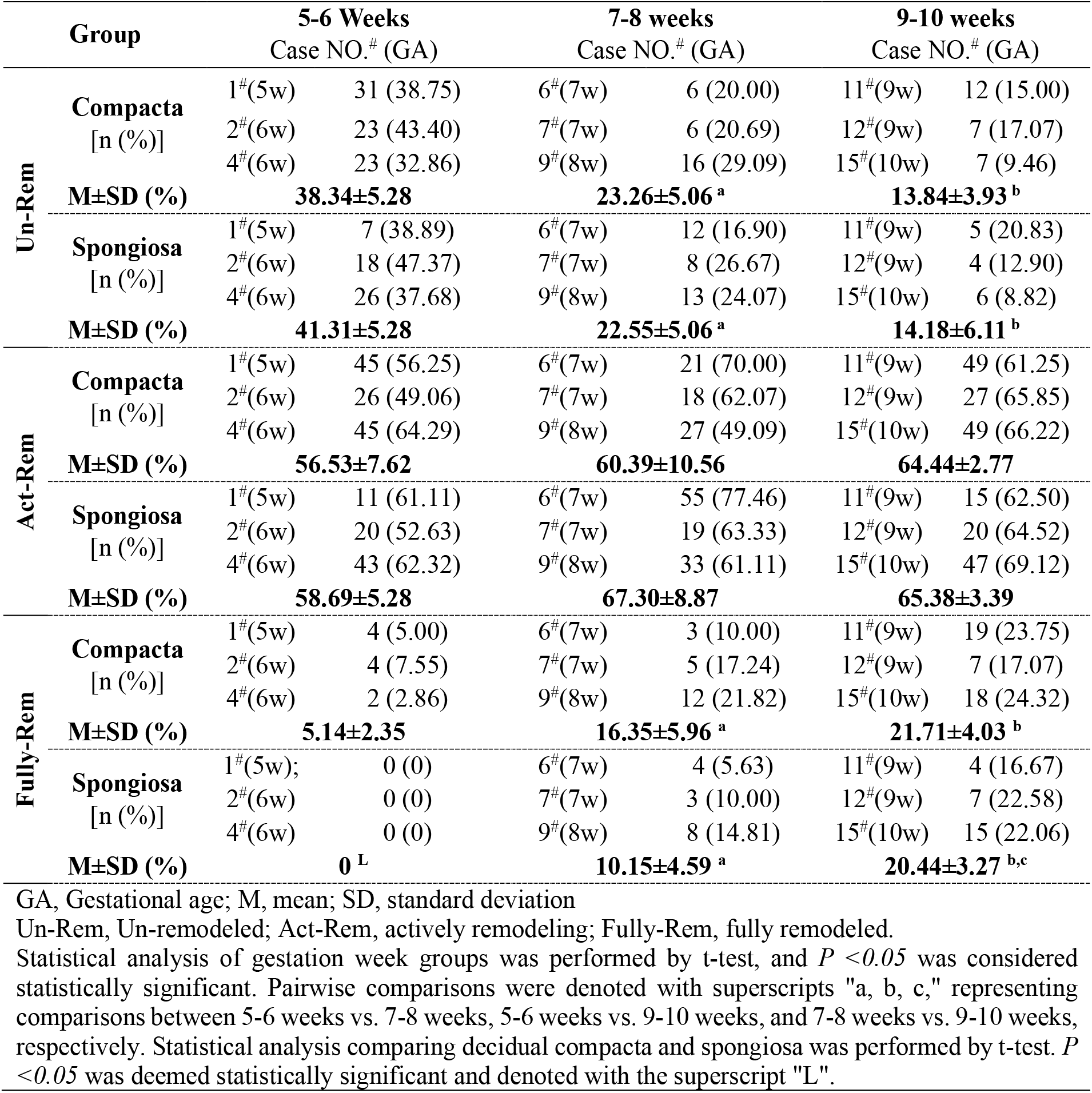
Spatiotemporal SAR in early human development.

### Spatiotemporal dynamics of EVT plugs

The formation of intraluminal HLA-G^+^/NCAM1^+^ endovascular EVT (enEVT) plugs is a pivotal event in SAR [15]. We further classified enEVT plugs into two subtypes based on their morphology and extent of lumen occupancy: 1) tight plugs composed of densely aggregated enEVTs occupying more than two-thirds of the vessel lumen and 2) loose plugs consisting of loosely associated enEVTs occupying less than two-thirds of the lumen (**Fig. 4A–B**). Using multiplex immunofluorescence imaging, we quantified these enEVT plug subtypes across gestational age groups (5-6, 7-8, and 9-10 weeks; n=3 per group) and by decidual zone (compacta vs. spongiosa), analyzing every 20th section from a total of 200 serial slices per specimen. As summed up in **Table 3**, enEVT plugs were exclusively detected in the compacta region at 5-6 weeks of gestation. From 7-10 weeks, plugs were observed in both compacta and spongiosa regions. Across all timepoints, the prevalence of SAs containing enEVT plugs was significantly higher in the compacta compared to the spongiosa, and this prevalence increased progressively with gestational age. Furthermore, tight plugs formed more frequently in the compacta decidua, while loose plugs were more commonly observed in the spongiosa (**Fig. 4C, Table 3**). These findings highlight a spatial and temporal regulation of enEVT plug formation during early pregnancy and support a compartment-specific role for trophoblast invasion in SAR progression.

**Table 3:** Spatiotemporal enEVT plug formation in early human development.

|  | Group | 5-6 Weeks |  | 7-8 weeks |  | 9-10 weeks |  |
| --- | --- | --- | --- | --- | --- | --- | --- |
|  |  | Case NO.# (GA) |  | Case NO.# (GA) |  | Case NO.# (GA) |  |
| Without Plug | **Compacta**<br>[n (%)] | 1<sup>#</sup>(5w) | 26 (92.85) | 6<sup>#</sup>(7w) | 13 (81.25) | 11<sup>#</sup>(9w) | 13 (65.00) |
|  |  | 3<sup>#</sup>(6w) | 27 (81.82) | 8<sup>#</sup>(7w) | 18 (66.67) | 12<sup>#</sup>(9w) | 20 (64.52) |
|  |  | 5<sup>#</sup>(6w) | 20 (83.33) | 10<sup>#</sup>(8w) | 14 (77.78) | 15<sup>#</sup>(10w) | 23 (67.65) |
|  | **M±SD (%)** | **86.00±5.98** |  | **75.23±7.62** |  | **65.72±1.69<sup>b</sup>** |  |
|  | **Spongiosa**<br>[n (%)] | 1<sup>#</sup>(5w) | 15 (100.00) | 6<sup>#</sup>(7w) | 21 (91.30) | 11<sup>#</sup>(9w) | 28 (80.00) |
|  |  | 3<sup>#</sup>(6w) | 9 (100.00) | 8<sup>#</sup>(7w) | 17 (80.95) | 12<sup>#</sup>(9w) | 18 (90.00) |
|  |  | 5<sup>#</sup>(6w) | 11 (100.00) | 10<sup>#</sup>(8w) | 12 (92.31) | 15<sup>#</sup>(10w) | 26 (89.66) |
|  | **M±SD (%)** | **100.00±0<sup>L</sup>** |  | **88.19±6.29<sup>a</sup>** |  | **86.55±5.68<sup>b,L</sup>** |  |
| Loose Plug | **Compacta**<br>[n (%)] | 1<sup>#</sup>(5w) | 0 (0) | 6<sup>#</sup>(7w) | 0 (0) | 11<sup>#</sup>(9w) | 2 (10.00) |
|  |  | 3<sup>#</sup>(6w) | 2 (6.06) | 8<sup>#</sup>(7w) | 4 (14.81) | 12<sup>#</sup>(9w) | 4 (12.90) |
|  |  | 5<sup>#</sup>(6w) | 1 (4.17) | 10<sup>#</sup>(8w) | 1 (5.56) | 15<sup>#</sup>(10w) | 4 (11.76) |
|  | **M±SD (%)** | **3.41±3.10** |  | **6.79±7.48<sup>a</sup>** |  | **11.55±1.46<sup>b</sup>** |  |
|  | **Spongiosa**<br>[n (%)] | 1<sup>#</sup>(5w) | 0 (0) | 6<sup>#</sup>(7w) | 2 (8.70) | 11<sup>#</sup>(9w) | 5 (14.29) |
|  |  | 3<sup>#</sup>(6w) | 0 (0) | 8<sup>#</sup>(7w) | 3 (14.29) | 12<sup>#</sup>(9w) | 2 (10.00) |
|  |  | 5<sup>#</sup>(6w) | 0 (0) | 10<sup>#</sup>(8w) | 1 (7.69) | 15<sup>#</sup>(10w) | 2 (6.90) |
|  | **M±SD (%)** | **0** |  | **10.23±3.56<sup>a</sup>** |  | **10.40±3.71<sup>b</sup>** |  |
| Tight Plug | **Compacta**<br>[n (%)] | 1<sup>#</sup>(5w) | 2 (7.14) | 6<sup>#</sup>(7w) | 3 (18.75) | 11<sup>#</sup>(9w) | 5 (25.00) |
|  |  | 3<sup>#</sup>(6w) | 4 (12.12) | 8<sup>#</sup>(7w) | 5 (18.52) | 12<sup>#</sup>(9w) | 7 (22.58) |
|  |  | 5<sup>#</sup>(6w) | 3 (12.50) | 10<sup>#</sup>(8w) | 3 (16.67) | 15<sup>#</sup>(10w) | 7 (20.59) |
|  | **M±SD (%)** | **10.59±2.99** |  | **17.98±1.14<sup>a</sup>** |  | **22.72±2.21<sup>b,c</sup>** |  |
|  | **Spongiosa**<br>[n (%)] | 1<sup>#</sup>(5w) | 0 (0) | 6<sup>#</sup>(7w) | 0 (0) | 11<sup>#</sup>(9w) | 2 (5.71) |
|  |  | 3<sup>#</sup>(6w) | 0 (0) | 8<sup>#</sup>(7w) | 1 (4.76) | 12<sup>#</sup>(9w) | 0 (0) |
|  |  | 5<sup>#</sup>(6w) | 0 (0) | 10<sup>#</sup>(8w) | 0 (0) | 15<sup>#</sup>(10w) | 1 (3.45) |
|  | **M±SD (%)** | **0<sup>L</sup>** |  | **1.59±2.75<sup>L</sup>** |  | **3.05±2.88<sup>L</sup>** |  |

GA, Gestational age; M, mean; SD, standard deviation.
Statistical analysis of gestation week groups was performed by t-test, and $P < 0.05$ was considered statistically significant. Pairwise comparisons were denoted with superscripts "a, b, c," representing comparisons between 5-6 weeks vs. 7-8 weeks, 5-6 weeks vs. 9-10 weeks, and 7-8 weeks vs. 9-10 weeks, respectively. Statistical analysis comparing decidual compacta and spongiosa was performed by t-test. $P < 0.05$ was deemed statistically significant and denoted with the superscript "L".

**Figure 4.**
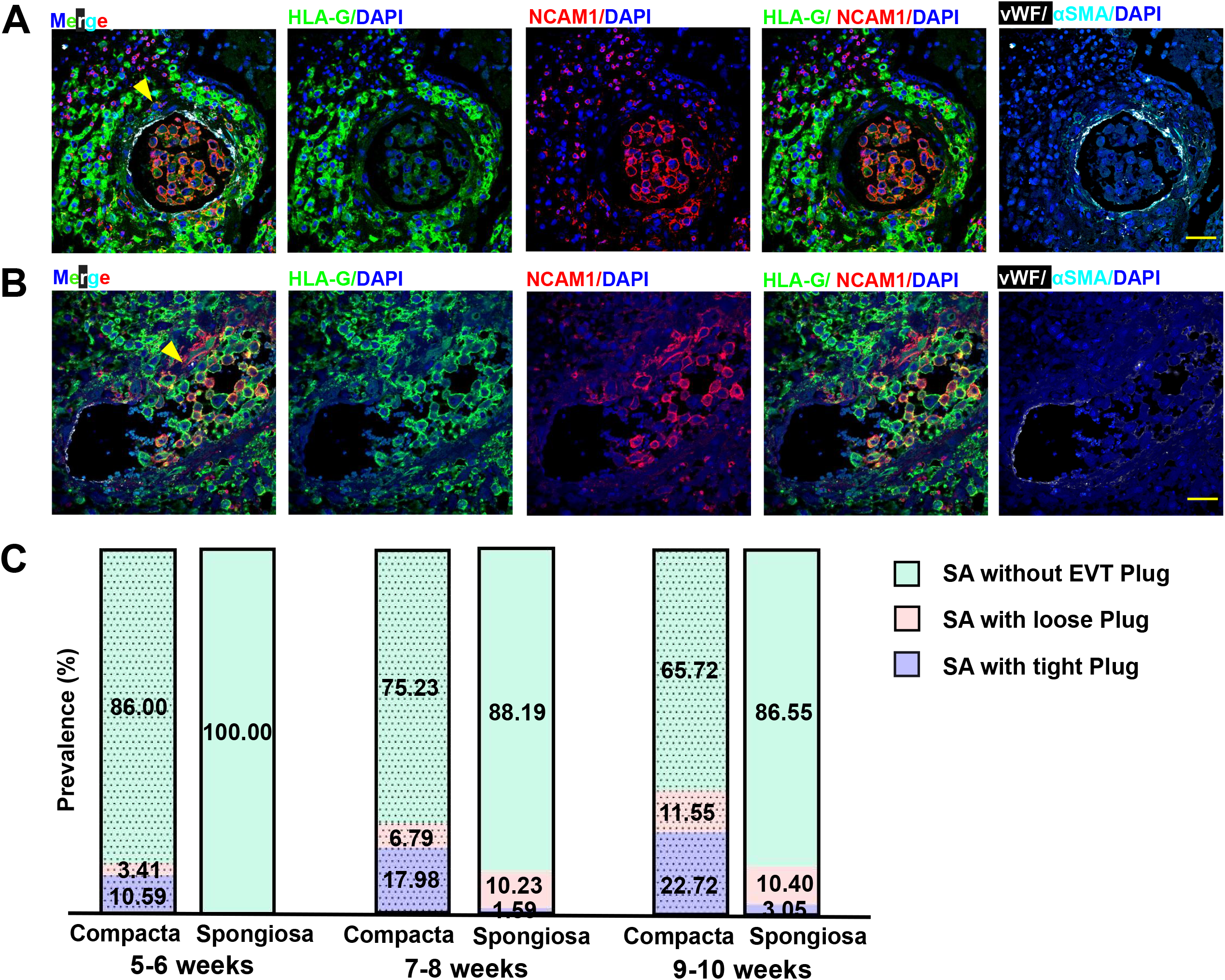
Dynamic changes in endovascular EVT (enEVT) plugs at the maternal–fetal interface during 5– 10 weeks of gestation. A-B. Representative multiplex immunofluorescence images of spiral arteries (SAs) in the decidua basalis showing different subtypes of enEVT plugs within the arterial lumen: (B) Tight plug in a GW7 specimen, characterized by densely packed enEVTs occupying >2/3 of the lumen. (C) Loose plug in a GW8 specimen, composed of loosely aggregated enEVTs occupying <2/3 of the lumen. Green: HLA-G (EVTs); Red: NCAM1 (decidual natural killer [dNK] cells); White: von Willebrand factor (vWF, endothelial cells); Cyan: α-smooth muscle actin (αSMA, smooth muscle cells); Blue: DAPI (nuclei). Yellow arrows indicate enEVT plugs. Scale bar: 50 μm. C. Bar graphs summarizing the mean prevalence of tight and loose enEVT plugs in the compacta and spongiosa layers of the decidua basalis across gestational ages. All SAs in each stained section were analyzed. Approximately 200 serial sections were prepared per case, and every 20th section was stained and quantified. Detailed statistics for enEVT plug prevalence by gestational week are provided in Table 3.

## DISCUSSION

Spiral artery remodeling is a critical event in hemochorial placentation and is conserved across rodents, nonhuman primates, and humans. The similarities in placental development among these species have facilitated the use of animal models to gain valuable insights into human SAR [34, 35]. However, significant interspecies differences in trophoblast differentiation and invasion, cell compositions, and cell-cell interactions at the maternal-fetal interface limit the direct applicability of these animal models to understanding physiological SAR in humans. To comprehensively elucidate physiological SAR during human placentation and investigate abnormal SAR linked to pregnancy complications such as miscarriages, fetal growth restriction, and the uniquely human pregnancy disorder preeclampsia, fresh human maternal-fetal interface specimens are indispensable [14, 36]. Currently, specimens obtained from elective abortion procedures between 5 and 10 weeks of gestation represent the primary source for cellular and molecular mechanistic studies of human placentation. However, obtaining high-quality specimens with an intact maternal-fetal interface remains challenging due to tissue fragmentation and dispersion caused by the surgical procedure. Variability in collection and processing protocols contributes to substantial differences in sample quality, which likely underlies the inconsistent or even contradictory observations of SAR dynamics reported across different studies [3, 11, 14, 15, 17-20, 23, 28, 32].

Endometrial stromal cells differentiate into specialized decidualized cells approximately 5-8 days post-fertilization. Decidualization initiates at the site of embryo implantation and subsequently forms the functional decidua basalis which contains the maternal-fetal interface. This process then extends throughout the uterine endometrium, resulting in the formation of the decidua capsularis and decidua parietalis. Our morphological and immunohistochemical analyses of aborted tissue specimens confirmed the presence of three distinct decidual tissue types corresponding to the capsular, parietal, and basal decidua. These detailed structural characterizations enabled us to develop a standardized protocol for isolating basal decidua specimens that retain an intact maternal-fetal interface from the heterogeneous mixture of aborted tissues. Moreover, we demonstrate that these isolated decidua basalis tissues are well-suited for quantifying the spatiotemporal dynamics of SAR. Utilizing multiplexed immunofluorescence microscopy with specific markers for the major cell types at the maternal-fetal interface, including EVTs, dNK cells, ECs and VSMCs, our analyses have established a robust and reproducible method to quantify physiological SAR dynamics from 5 to 10 weeks of gestation using fresh human maternal-fetal interface specimens derived from aborted materials.

Notably, we observed that SAR initiates in SAs located within the decidua compacta as early as week 5 of gestation, characterized by infiltration of NCAM1^+^ dNK cells and migration of NCAM1^−^/HLA-G^+^ iEVTs into the SA’s medial layer (**Fig. 3A**). Luminal NCAM1^+^/HLA-G^+^ enEVTs first appear in actively remodeling SAs during weeks 5-6, coinciding with the formation of tight enEVT plugs. As gestation progresses, these tight plugs transition into loose plugs, with a progressive replacement of EC layer by enEVTs. These observations align with prior studies that describe SAR as primarily an intravascular process[8]. Furthermore, NCAM1^+^ dNK cells and NCAM1^−^/HLA-G^+^ iEVTs initially localize to the interstitial space surrounding un-remodeled SAs and subsequently spread to actively and fully remodeled SAs as gestation advances (**Fig. 3**). Given that dNK cells can directly disrupt the arterial muscular lining and indirectly facilitate iEVT invasion into SAs [37], our data support the notion in which SAR begins outside the vessels with recruitment of dNK cells. Additionally, approximately 56% of SAs undergo active remodeling in both the compacta and spongiosa decidua during weeks 5-6 (**Fig. 3A & B**), temporally coinciding with CTB differentiation into EVTs around day 21 post-fertilization [38]. Our findings on SAR progression in compacta versus spongiosa decidua across weeks 5-6, 7-8, and 9-10 (**Fig. 3C-D**) are consistent with previous reports describing SA remodeling initiation at the implantation site (compacta decidua) followed by distal progression toward the spongiosa decidua [15]. However, we also found that the lumen areas of un-remodeled SAs gradually increase from weeks 5 to 10, suggesting a novel enEVT-independent pathway for remodeling. This may involve dedifferentiation of VSMCs through interactions with decidual stromal and immune cells [32].

We also observed that tight enEVT plugs form in fewer than 10% of SAs at week 5 of gestation, while loose enEVT plugs begin to appear gradually after week 7. Subsequently, the proportion of SAs containing loose plugs progressively increases in the decidua compacta as gestation advances (**Fig. 4, Table 3**). These findings indicate a critical role of the formation of enEVT plugs and their temporal transition from tight to loose plug morphology in SAR. Specifically, tight plugs formed during weeks 5-6 obstruct blood flow, creating a hypoxic microenvironment that promotes angiogenesis and vessel expansion at this early stage [20, 23]. The loosening of these plugs, coinciding with a significantly higher proportion of fully remodeled SAs at weeks 9-10, signifies the completion of SAR and the establishment of blood flow into the intervillous space. Despite these insights, the mechanisms regulating the formation and dissolution of enEVT plugs throughout SAR, as well as the process by which enEVTs replace ECs remain to be elucidated. Notably, vessel lumen areas are smallest in un-remodeled SAs, increase during active remodeling and reach their largest size in fully remodeled vessels. This underscores that enEVT-mediated intraluminal remodeling not only enlarges vessel diameter but also reconstructs the vessel wall architecture.

While our current data are consistent with previous studies showing that endovascular enEVTs replace ECs in fully remodeled SAs [8], the molecular characteristics of enEVT plugs remain poorly understood. Our analysis reveals that enEVT plugs consist of heterogeneous cell populations, exhibiting diverse expression patterns of HLA-G and NCAM1 [22]. This heterogeneity aligns with our recent findings that enEVTs possess immune regulatory functions, including secretion of transforming growth factor-β (TGF-β) [21]. The expression of NCAM1 suggests that enEVTs may acquire NK cell-like properties [27], potentially equipping them with the functional capacity to mediate intraluminal vascular remodeling. This hypothesis is further supported by emerging spatial multiomics studies highlighting the functional plasticity of enEVTs [16, 29]. Moreover, the acquisition of immune regulatory properties may enable enEVTs to contribute to the establishment of immune tolerance at the maternal-fetal interface. However, the precise timing and mechanisms by which enEVTs acquire these immune functions remain to be elucidated.

Multiplex immunofluorescence microscopy enables simultaneous visualization of multiple maternal and fetal cell populations within serial sections from the same specimen, as well as across samples from different gestational ages. This approach yields novel insights into the cellular composition and spatiotemporal cell-cell interactions that drive SAR. While recent studies using single-cell and spatial transcriptomics have significantly advanced our understanding of the cellular landscape at the human maternal–fetal interface [24], these analyses are typically based on archived samples, which often lack precise gestational age resolution. In contrast, applying such high-dimensional analyses to fresh specimens collected using our standardized protocol could provide more continuous and developmentally informative data. This would allow for a finer-resolution mapping of the dynamic cellular and molecular changes occurring throughout early gestation.

It is estimated that up to 200 SAs open into the intervillous space to supply maternal blood during human pregnancy [39], although this number can vary substantially depending on gestational age [40]. In this study, we were able to quantify mean vessel lumen areas; however, we were unable to determine the total number of SAs contributing to the intervillous circulation during weeks 5-10 of gestation. This limitation arises from the nature of elective abortion procedures, which often disrupt the structural integrity of the maternal-fetal interface. As a result, it is challenging to obtain intact tissue specimens suitable for imaging for accurately counting the total number of SAs connected to the intervillous space.

In this study, we analyzed samples from a total of 15 patients and grouped the analyzed cases biweekly into three groups, i.e., 5-6, 7-8, and 9-10 weeks of gestation, for statistical analysis of SAR according to gestational age and depth of decidua (compacta vs. spongiosa). While the relatively small sample size (n=3/group) may limit the generalizability of our findings, the analytical depth of our approach strengthens the reliability of our findings and suffices the validation of our comprehensive methodology. Specifically, we analyzed at least 10 sections from every 20th slice across 200 serial sections per sample. This comprehensive sampling strategy enables accurate assessment of spatial changes in SAR and supports robust quantitative analysis of SAR progression during early gestation, although increasing sample size per gestation age will help solidify our current fundings.

Aberrant or impaired SAR is implicated in the pathogenesis of nearly all major human pregnancy complications [12], highlighting the need to investigate the mechanisms underlying physiological SAR. Herein we analyzed samples obtained from pregnancies without known complications, thereby providing insight into normal SAR processes. Using these well-characterized samples, we previously uncovered a novel fate of VSMCs during SAR; specifically, approximately 5% of VSMCs undergo dedifferentiation and transform into dNK cells [32]. Importantly, the same protocol has also been successfully applied to identify pathological maternal-fetal interface specimens from patients with recurrent miscarriages; these analyses revealed impaired immunoregulatory functions of enEVTs and insufficient SAR in patients with recurrent miscarriages [21, 22]. These findings demonstrate the utility of our protocol not only for studying physiological SAR but also for investigating pathological SAR associated with abnormal placentation and defective uteroplacental vascular remodeling. Thus, this approach can be extended to study SAR in pregnancy complications such as preeclampsia and fetal growth restriction, with potential applications in biomarker discovery and therapeutic target identification.

Altogether, our study establishes a standardized and reproducible protocol for classifying and isolating maternal-fetal interface tissues from heterogeneous aborted materials, enabling mechanistic investigations into early human development. Through multiplex immunofluorescence imaging with cell type–specific markers, we demonstrate that basal decidua tissues collected from gestational weeks 5 to 10 are of high quality and well-suited for comprehensive analyses of SAR. These specimens offer valuable insights into the morphological, structural, cellular, and molecular dynamics of physiological SAR during early human placentation, supporting the adoption of our methodology in future studies of abnormal SAR and related pregnancy disorders.

## Acknowledgement

The authors thank all patients for participating in this study. We appreciate Ms. Shiwen Li, Xili Zhu, and Yue Wang at the Institute of Zoology of the Chinese Academy of Sciences for technical support in Confocal microscopy.

## Author Contributions

Ye S, Chen DB, and Wang YL designed the study; Qi L, Yu D and Wang X identified and consent patients; Ye S, Ma Y, Li W, and Qi L processed samples, carried out experiments, and collected data; Fang X and Yu X collected data; Ye SL, Chen DB, Wang YL analyzed data; Ye SL drafted the paper which was finalized by Chen DB and Wang YL.

## Data Availability

Original data presented in the study are included in the article and Supplementary Data. Further inquiries about the study and samples can be directed at the corresponding authors.

